# Dietary and Reproductive Traits of Two Brachyuran Crab Species

**DOI:** 10.1101/2022.04.08.487630

**Authors:** Ben Potter, Zachary J. Cannizzo, Blaine D. Griffen

**Affiliations:** Biology Department, Brigham Young University, Provo, UT 84604; National Oceanic and Atmospheric Administration Office of National Marine Sanctuaries − National Marine Protected Areas Center, Silver Spring, MD 20910, USA

**Keywords:** *Aratus pisonii*, claw size, diet, functional traits, *Hemigrapsus sanguineus*, herbivory, fecundity

## Abstract

Many animals have flexible morphological traits that allow them to succeed in differing circumstances with differing diets available to them. For brachyuran crabs, claw height and gut size are diet-specific and largely reflect foraging strategies, while abdomen width reflects relative levels of fecundity. However, the link between claw size and diet has largely been documented only for primarily carnivorous crabs, while the link between diet and fecundity is strong in herbivorous crabs. We sought to determine the nature of the intraspecific relationship between claw size, dietary habits, and fecundity for two primarily herbivorous crab species, *Hemigrapsus sanguineus* and *Aratus pisonii*. Specifically, we examined whether claw size and/or abdomen width can be used as reliable measures of individual diet strategy. To test these hypotheses, we collected crabs and measured the dimensions of their claws, abdomens, and guts. By comparing these dimensions for each individual, we found that strongly predictive relationships do not exist between these traits for the primarily herbivorous species in our study. Thus, identifying external morphological features that can be used to assess diets of primarily herbivorous crabs remains elusive.

## Introduction

A long-standing goal of ecology is to predict how populations and systems will be affected by future changes to their habitats and resources, changes that continue to become more frequent due to increasing human pressures and environmental change. Making these predictions can be difficult when relying only on past and current population trends, so it has become increasingly common to use individual-based ecology when making predictions. Individual-based ecology focuses on fitness-based decisions of individuals within a population, allowing the development of broader population and ecosystem predictions based on individual traits and processes (Stillman et al., 2015).

Functional traits (i.e., individual level morphological or physiological traits that improve fitness) are useful tools in understanding the complexity of predator-prey relationships (Schmitz, 2017). These traits are often highly flexible, so their expression must be considered within the environmental context of the individual. The factors influencing this context include changes in resource quality and consumer pressure (Schmitz et al., 2015). These environmental influences result in differences in functional trait expression within a species due to geographic distribution (Smith, 2004). Because of the variability of functional traits, they represent a key aspect of understanding the driving influences of predator-prey relationships and their role in ecosystem functioning (Schmitz, 2017). By understanding how functional traits of an individual differ in response to resource availability, we can better predict a species’ ability to adapt to changes in the future.

Crabs are an ideal group for considering the role of functional traits in organisms’ responses to changes in resource availability because they have measurable physical traits that can be used to quantify their success as consumers. Individual diet variation is a crucial determinant of reproductive effort in crabs (Griffen, 2014; Riley et al., 2014; Griffen and Norelli, 2015; Griffen and Riley, 2015; Belgrad and Griffen, 2016; Gül and Griffen, 2020). As in other species, diet can be assessed in crabs using standard methods, such as stable isotope analysis or gut content analysis. However, stable isotopes are expensive and time consuming when conducted on large numbers of individuals, and diet identification using gut content analysis is difficult in crabs given their propensity for shredding and masticating their food. In addition, gut content analysis in crabs requires sacrificing animals, and can therefore only be measured a single time per individual.

In carnivorous crabs, well-known proxies exist for assessing diet quickly, cheaply, and nonlethally. For instance, claw size is highly flexible for many crab species, varying in response to resource quality and availability. Among the European green crab *Carcinus maenas*, captive individuals that were fed snails with harder shells developed larger claws than those fed snails with softer shells (Edgell & Rochette, 2009). Smith and Palmer (1994) similarly demonstrated that crabs raised on shelled prey developed larger and stronger claws than crabs raised on unshelled prey of equivalent nutritional value. These findings show that this trait is variable within an individual’s lifetime, rather than just on an evolutionary timescale. Observational studies have shown the same trend between claw size and shell hardness of the crab’s prey across geographical regions and habitats (Smith, 2004; Silva et al., 2010). Crabs living in regions with harder-shelled prey have greater crushing strength than crabs in regions with softer-shelled prey (Smith, 2004). Of the various dimensions of crab claws, claw height is the best predictor of strength and crushing ability (Smith, 2004; Wilcox & Rochette, 2015). The increase in claw height is presumably the result of the crabs’ necessity for greater crushing strength to break through the harder shells of their prey and feed on the soft interior. While this trend has been repeatedly shown in species that are primarily carnivorous, generalist species and species that are primarily herbivorous have been underrepresented in past claw height studies.

Abdomen width is another morphological feature of brachyuran crabs that acts as a functional trait, in this case as a good predictor of fecundity, and therefore of fitness. Studies on multiple crab species have demonstrated strong positive relationships between fecundity and abdomen width (Rodrigues et al., 2011; Ali & Al-Maliky, 2017; Litulo, 2004). Abdomen width can therefore be used as a measure of relative fecundity between individuals of the same species because individuals with greater abdomen widths generally have larger clutch sizes. As noted above, reproductive effort in crabs is highly dependent on diet. To our knowledge, no studies have examined changes in abdomen width with individual diet to determine whether abdomen width could be used as a diet predictor in addition to its role in predicting fecundity.

Yet another functional trait, gut size is a strong predictor of diet in brachyuran crabs (as well as in many other species) and can reliably be used to assess diet difference both between individuals and across sites. In a study of 15 different species of brachyuran crabs, gut size had a strong inverse relationship with the quality of the diet of individual crabs, both within and across species. Specifically, gut size served as a strong predictor (R^2^ = 0.80 after accounting for phylogenetic relationships among species) of the percent of herbivory in the crab’s diet, with greater herbivory leading to a larger gut (Griffen & Mosblack, 2011). This morphological difference is most likely the result of the low digestibility and nutrient content, especially nitrogen, of many plant materials. Due to the relatively low nutrient content, a greater volume of plant material is needed to meet the same metabolic requirements that can be met with a smaller volume of animal material (i.e., compensatory feeding), which results in a larger gut over time for more herbivorous individuals. Previous studies demonstrate that gut width is a reliable predictor of diet in individual crabs (Griffen & Mosblack, 2011; Griffen et al., 2012), as well as at the population level by comparing across sites with different levels of food availability (Cannizzo et al., 2017, 2020; Griffen et al., 2020a).

Our study considers two brachyuran crab species, the Asian shore crab *Hemigrapsus sanguineus* and the mangrove tree crab *Aratus pisonii*. Both species are primarily herbivorous (Griffen et al., 2020a; Griffen et al., 2020b). *H. sanguineus* has an omnivorous diet that can include macroalgae and detritus, as well as amphipods, polychaetes, and juvenile shelled prey, such as barnacles, bivalves, and gastropods (Brousseau & Goldberg, 2007). They primarily consume macroalgae but will consume animal material when it is present (Brousseau & Baglivo, 2005). *A. pisonii* is also primarily herbivorous, with most of its diet consisting of the leaves of mangrove trees, especially the red mangrove, *Rhizophora mangle* (Riley et al., 2015 and references therein). In addition, they have been documented feeding on insects, gastropods, dead fish, and other crabs (Beever et al., 1979; Erickson et al., 2008). Like *H. sanguineus, A. pisonii* has been described as an opportunistic omnivore, feeding on animal material preferentially when it is available, while plant material still makes up the bulk of its diet (Erickson et al., 2008). Finally, gut size is a reliable predictor of diets of individual crabs for both of these species (Griffen & Mosblack, 2011; Griffen et al., 2012; Cannizzo et al., 2017, 2020; Griffen et al., 2020a).

In the present study, we compare intraspecific variation and correlations between claw height and gut size, and between abdomen width and gut size for both *H. sanguineus* and *A. pisonii* to determine whether claw height and/or abdomen width can be used as reliable proxies for diet strategy in individuals of these two primarily herbivorous species. Fecundity generally increases with the amount of animal tissue included in the diet for crab species in general (Griffen, 2014; Griffen & Norelli, 2015; Griffen & Riley, 2015; Gül & Griffen, 2020), and for these species in particular (Riley et al., 2014; Griffen et al., 2020). If external morphology is a reliable predictor of diet-driven fitness, we therefore expect claw height (i.e., strength) and abdomen width to each be inversely correlated with gut width. By describing these relationships within these two primarily herbivorous species, we hope to provide tools to better predict how these species will react to future changes in resource availability.

## Methods

### Sample sites & crab collection

We sampled populations of *H. sanguineus* and *A. pisonii* from sites along the East coast of the United States. We only sampled female crabs because we used abdomen width as a metric of fecundity, and this relationship is only present in females. Additionally, measuring female claws can help avoid confounding factors, such as claw size selected for non-feeding purposes, such as for defense and mating, that are more common with males (Yamada & Boulding, 1998).

*H. sanguineus* samples were gathered in August 2019 from two different sites in Rye, New Hampshire: Fort Stark and Odiorne Point State Parks. Both sites are comprised of boulder fields, and they contain the same group of intertidal algal and animal species. However, Odiorne Point has a much greater density of *H. sanguineus* than Fort Stark, which has led to a decreased abundance of prey species at Odiorne Point (Griffen et al., 2021). Because Fort Stark has more animal prey and shelled prey available per capita than Odiorne, we expect differences in diet to exist between the two populations. *H. sanguineus* samples were collected haphazardly at dawn by overturning boulders in the intertidal region. Only mature crabs (exceeding 12.1 mm carapace width) were collected. Upon collection, the samples were frozen and stored on dry ice at −80° C until dissection. Following dissection, the samples were dried at 65° C (see Griffen et al., 2020a for a complete description of sampling methods and for a more complete description of these two sites).

Individual *A. pisonii* were sampled from four sites in Florida along the Atlantic Intracoastal Waterway. Each of the four sites were fringing mangrove habitats that had similar levels of resource availability, but that differed in relative human disturbance. We rotated sampling between the four sites across sampling dates to avoid depleting any one population. The samples were originally collected for a study that explored reproduction (Griffen et al., 2020b), so they were collected before each spring tide from March-October in order to encompass the time leading up to and during their reproductive season. During sampling, the first 20 female crabs encountered during each sampling date were collected by hand and immediately frozen with dry ice and stored at −80° C until dissection. Following dissection, the samples were dried at 65° C.

### Morphological Measurements

To compare claw size and fecundity with herbivory levels, we measured dimensions of the claws, abdomen, and cardiac stomach (hereafter ‘gut’) for each crab. After the samples had been dissected and dried, we measured the gut width along the anterior portion of the cardiac stomach to the nearest 0.1 mm using Vernier calipers, using methods described in Griffen and Mosblack (2011). We used the same calipers to measure the abdomen at its widest point. We then removed the claws from the crab body and measured the claw height (Figure 1). For the *H. sanguineus* samples, only one claw was measured (the left-hand claw, if present) because this species is homochelous. Both claws were measured for *A. pisonii* because this species is heterochelous.

**Figure 1.**
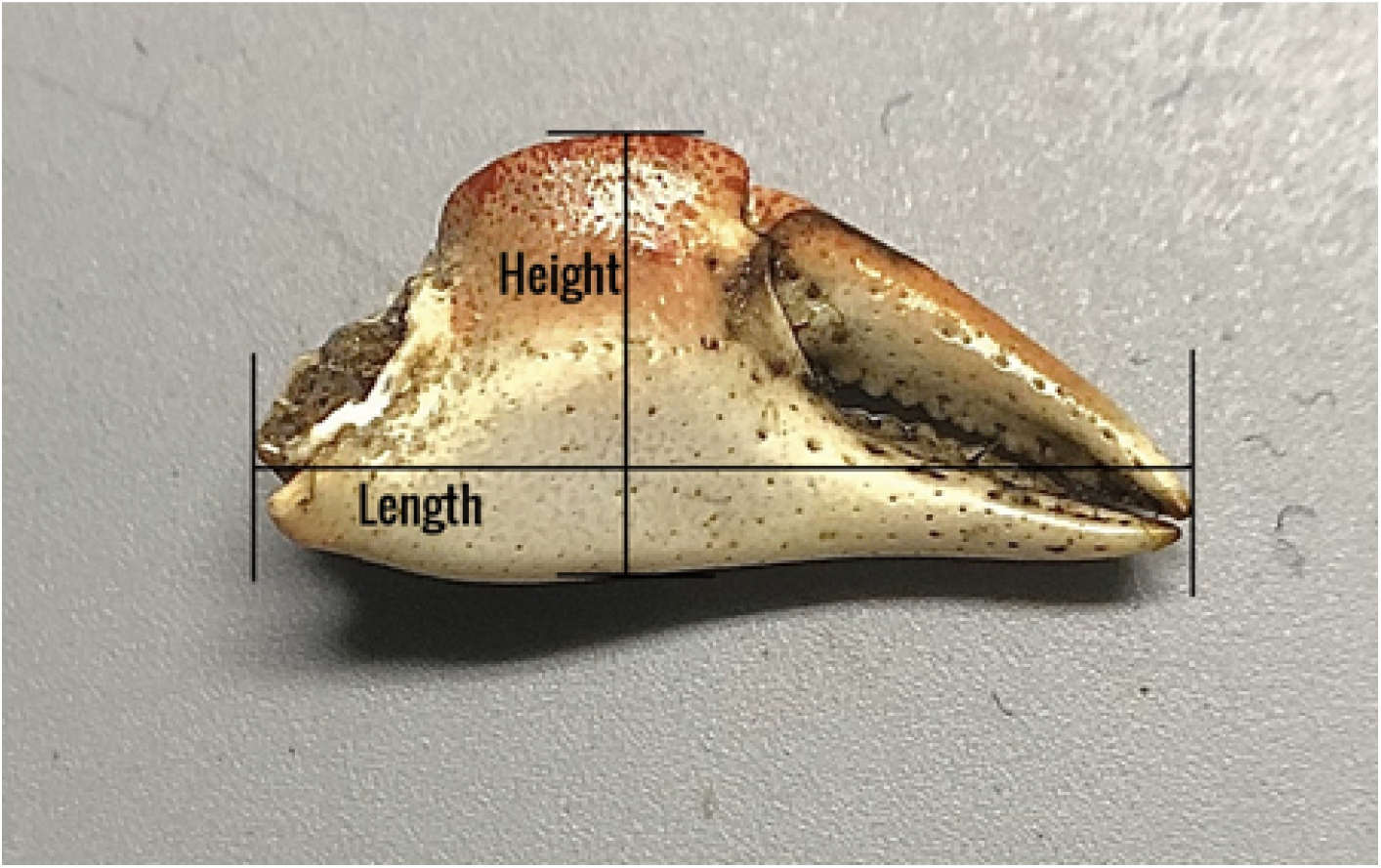
Dimensions for claw measurement. Here we focus only on differences in claw height, a proxy for claw strength in crabs.

### Analysis

We examined the relationship between claw height (i.e., a proxy for strength) and gut width (i.e., a proxy for diet) between crabs. We also examined the relationship between abdomen width (i.e., a proxy for reproductive potential) and gut width. Each of these measurements is correlated with body size due to allometric relationships. We therefore controlled for body size in our analysis by regressing each of these variables against carapace width using nonlinear regression, and then using the residuals from these regressions in linear models. We analyzed the two species separately, but using the same approach. For both species, we used residual gut width as the response variable, and residual abdomen width and residual claw height as continuous predictor variables, and collection site as a categorical predictor variable. For *A. pisonii*, residual claw height and residual abdomen width were positively correlated (major claw: R^2^ = 0.82, minor claw R^2^ = 0.20). To avoid multicollinearity, we therefore used residual claw height after controlling for both body size and abdomen width. For *A. pisonii*, significant differences in claw size across collection sites were compared using Tukey’s HSD pairwise *post hoc* comparisons. We only analyzed the single claw for *H. sanguineus* (using the left claw, unless it was missing), while for *A. pisonii* we separately analyzed both the major and minor claws.

## Results

### Hemigrapsus sanguineus

Our analyses included a total of 188 adult female crabs, 93 collected from Ft. Stark, and 95 collected from Odiorne Point State Park. After accounting for differences associated with crab body size by using residuals, we found that gut width increased by 0.13 ± 0.03 mm for every 1-mm increase in abdomen width (*t* = 3.62, *P* = 0.0004, Fig. 2A), but that gut width was not related to claw height (*t* < −0.18, *P* = 0.86, Fig. 2B), and was larger by 0.11 mm in crabs collected from Odiorne Point than in crabs collected at Fort Stark (*t* = 1.91, *P* = 0.057, Fig. 2C).

**Figure 2.**
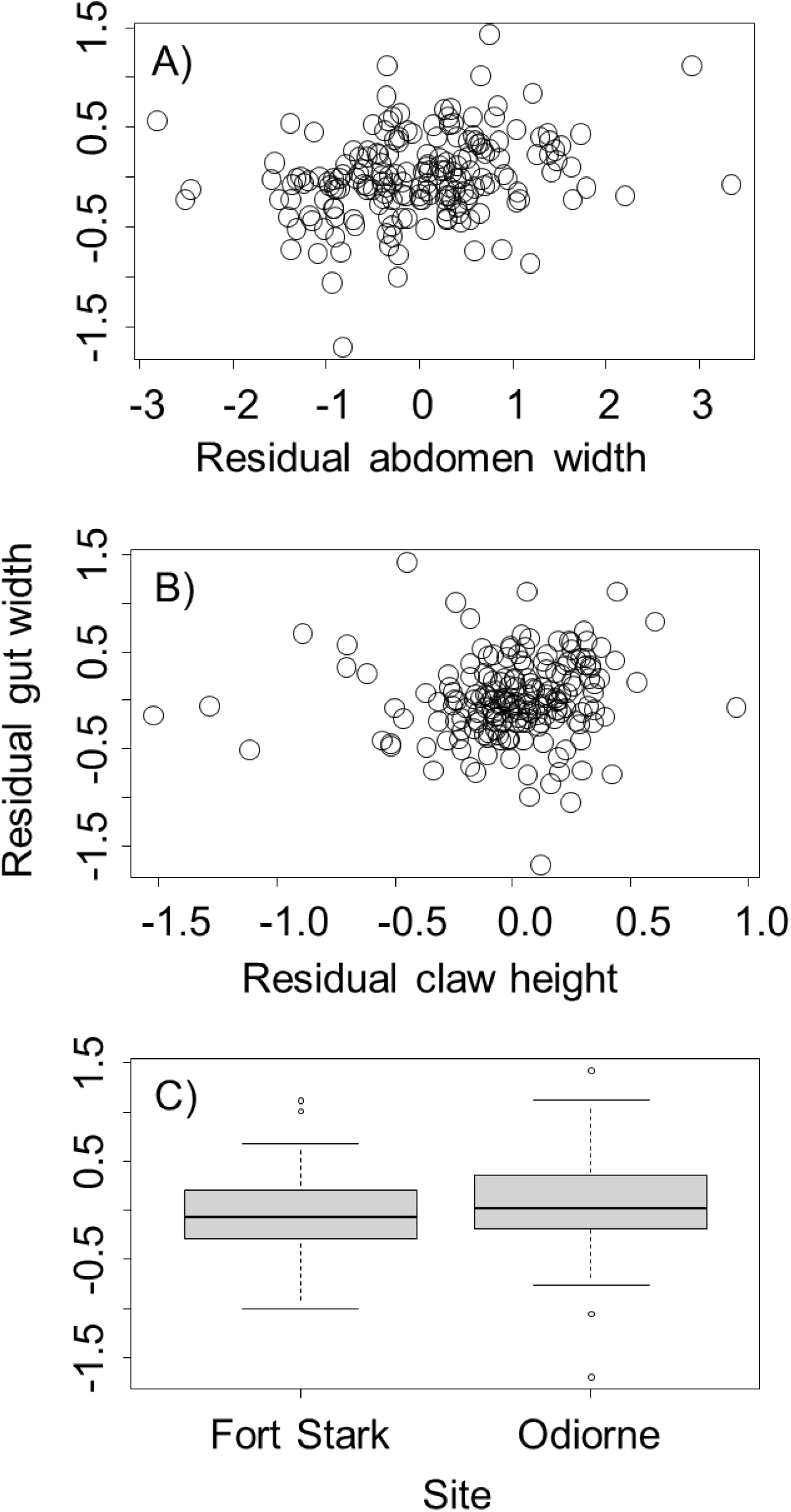
Relationships between residual gut width for *Hemigrapsus sanguineus* and residual abdomen width (A), residual claw height (B), and collection site (C). All residuals reflect sizes after accounting for differences associated with crab body size. Box plots in part C show the median value (heavy black line), the first to third quartile of the data (box), 95% of the data (whiskers), and outliers that fall outside this range (circles).

### Aratus pisonii

We compared the same metrics for 241 *A. pisonii* samples, examining both the major and minor claws. When using the major claw for analysis, after accounting for differences associated with crab body size by using residuals, we found that gut width increased by 0.04 ± 0.01 mm for every 1-mm increase in abdomen width (*t* = 2.73, *P* = 0.007, Fig. 3A), but was not related to the height of the major claw (*t* = 0.62, *P* = 0.53, Fig. 3B). We also found differences in gut width between sites. Specifically, crabs collected from Oslo had smaller guts than crabs collected from Round Island (*t* = 2.66, *P* = 0.042, Fig. 3C) and from Pepper Park (*t* = 2.65, *P* = 0.043, Fig. 3C). there were no other differences in gut size between sites (*P* > 0.05).

**Figure 3.**
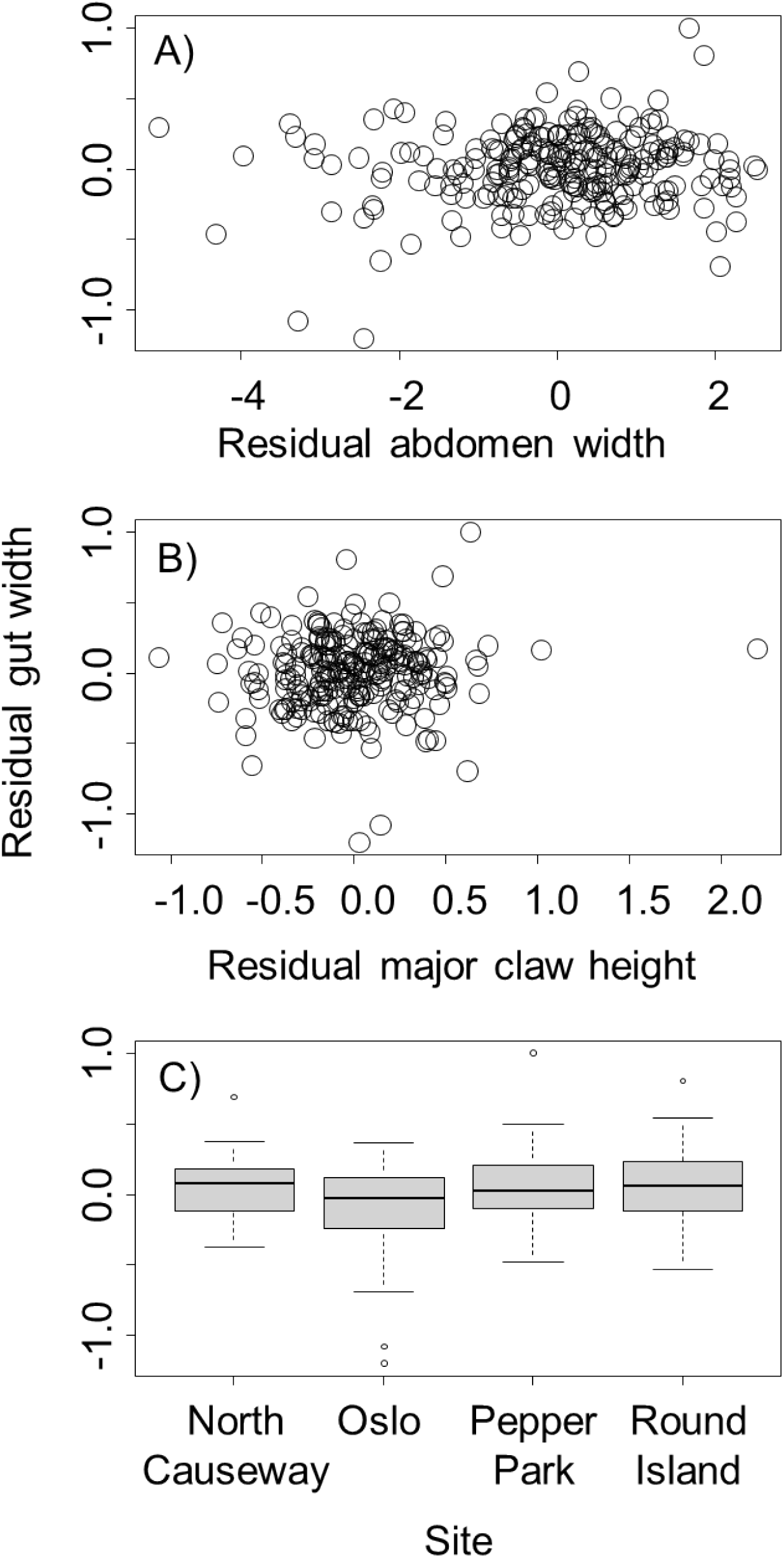
Relationship between residual gut width for *Aratus pisonii* and residual abdomen width (A), major claw residual height (B), and collection site (C). Residual gut width and abdomen width reflect sizes after accounting for differences associated with crab body size, which residual claw height reflects heights after accounting for correlations with both body size and abdomen width. Boxplots are as described in the caption for Fig. 2.

Patterns when the minor claw was used in the analysis were somewhat different. Specifically, we found that the gut width increased by 0.08 ± 0.03 mm for each 1-mm increase in minor claw height (*t* = 2.56, *P* = 0.011, Fig. 4A). However, there was no relationship between gut width and abdomen width (*t* = −1.31, *P* = 0.19, Fig. 4B). Finally, for completeness we also included collection site in this analysis, and it returned the same results qualitatively as the analysis with major claw height. Specifically, crabs collected from Oslo had smaller guts than crabs collected from Round Island (*t* = 2.95, *P* = 0.019, Fig. 4C) and Pepper Park (*t* = 2.96, *P* = 0.018, Fig. 4C). No other differences in gut size were observed between crabs collected from other sites (*P* > 0.05).

**Figure 4.**
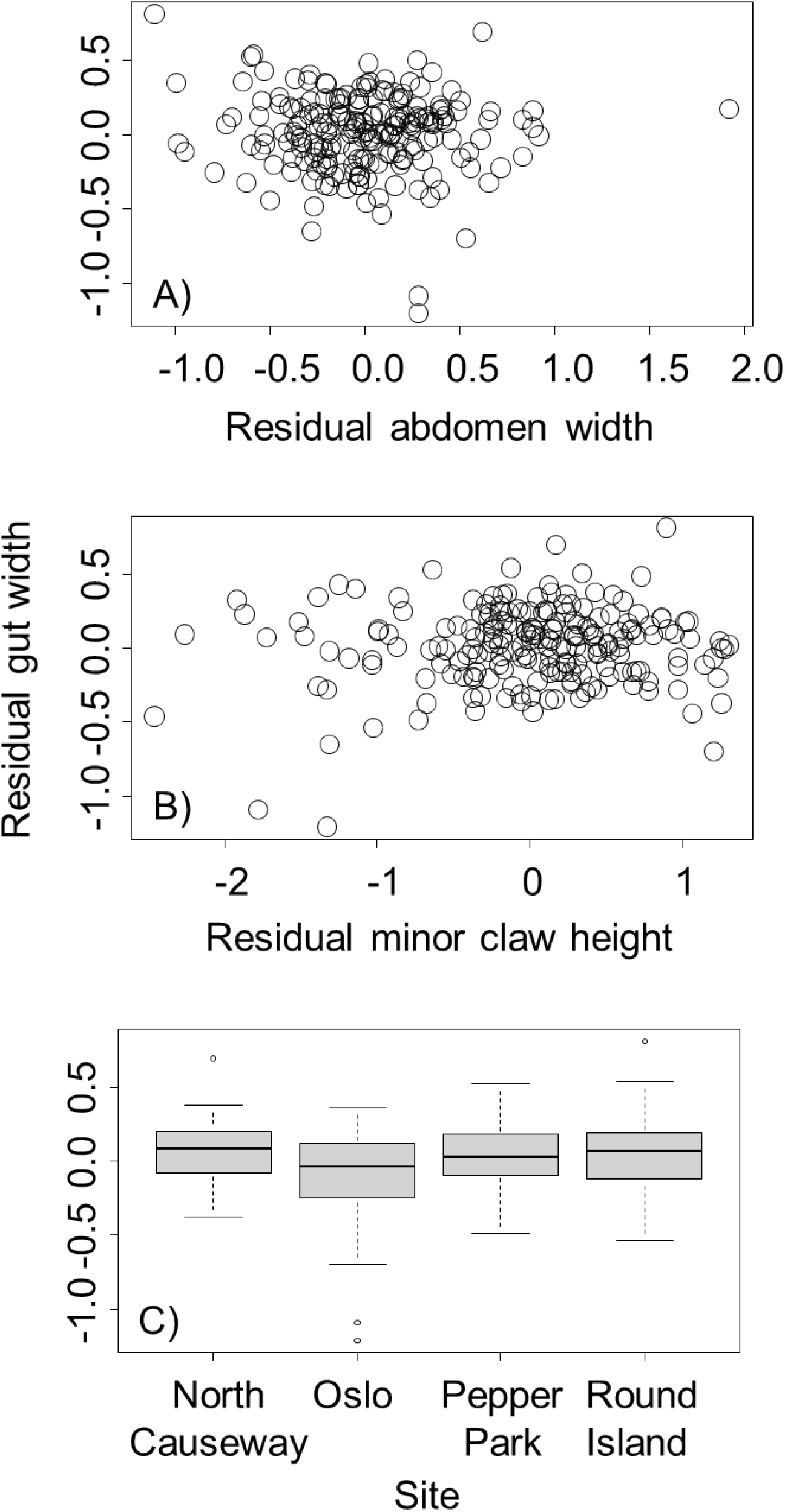
Relationship between residual gut width for *Aratus pisonii* and residual abdomen width (A), minor claw residual height (B), and collection site (C). Residual gut width and abdomen width reflect sizes after accounting for differences associated with crab body size, which residual claw height reflects heights after accounting for correlations with both body size and abdomen width. Boxplots are as described in the caption for Fig. 2.

## Discussion

In this study, we examined morphometric data of two primarily herbivorous species of brachyuran crabs to determine whether claw height and/or abdomen width can be used as reliable proxies for diet strategy in individuals of these species. We used gut size as a measure of the relative percent herbivory in the individual’s diet. Our results indicate that weak relationships exist between abdomen width and gut size for both species, but these relationships are likely not strong enough to use abdomen width as a reliable proxy for individual diet strategy. Further, the weak positive relationship between gut width and abdomen width is opposite to the negative correlation that we expected. No relationship was found between claw height and gut size for either species, other than a small correlation with the minor claw of *A. pisonii*. Thus, neither of these external morphological features provide a good proxy for individual diet strategy in these two species.

The results for *H. sanguineus* and *A. pisonii* indicate that claw height was not related to percent herbivory of the crab’s diet for either species. Among *H. sanguineus*, those collected from the site with less shelled and animal prey available (Odiorne) had greater claw heights than those collected from the site with more of these prey available (Fort Stark), counter to expectations based on studies of carnivorous species (Smith & Palmer, 1994; Smith, 2004). The differences between sites that existed among the *A. pisonii* samples were minimal, and they were likely unrelated to percent herbivory because all the sites presumably had similar levels of animal resource availability. These results show that the relationship between claw height and herbivory for these species does not follow the trends found in other crab species that are primarily carnivorous. This may be expected, as herbivorous species do not use their claws to break into hard-shelled prey. Thus, despite the ready consumption of animal prey when available, breaking into hard-shelled prey is not a measurable driver of claw size in *H. sanguineus* or *A. pisonii*. These findings are consistent with an experimental study performed by Yamada & Boulding (1998), which found that claw dimensions were strong predictors of closing force for two species that specialized in shell breaking (*Lophopenopeus bellus* and *Cancer productus*), but not for a generalist species (*Hemigrapsus nudus*) that does not specialize on hard-shelled prey. Our results suggest that *H. sanguineus* and *A. pisonii* may meet metabolic nitrogen requirements by scavenging dead animals (Wolcott, 1978), and/or consuming relatively soft animals (worms, gammarids, etc.).

The abdomen width also did not provide a clear proxy for assessing diet strategy. As explained in this Introduction, we anticipated a negative relationship between abdomen width and gut width. Instead, we observed weak positive relationships between abdomen width and gut width for both species. This surprising result may potentially be explained by anatomical constraints that disproportionately affect crab species that are primarily herbivorous. We have previously demonstrated that residual gut size increases with percent herbivory in these species (Griffen and Mosblack, 2011; Cannizzo et al., 2017, 2020), but this relationship appears to break down at very high levels of herbivory (>60%) (Quezada-Villa et al., *In review*). This may result from the limited space inside the carapace and the tradeoff in space use amongst the gut and other organs (gills, hepatopancreas, gonads, etc.). Because both of the species in our study are highly herbivorous, they are both likely affected by this constraint, thus preventing the expected negative relationship between gut size and abdomen width.

In conclusion, multiple studies demonstrate a clear relationships between individual diet strategy and claw size among primarily carnivorous crab species (Edgell & Rochette, 2009; Smith & Palmer, 1994), but our study demonstrates that neither claw height nor abdomen width can reliably be used to estimate individual diet strategy for these two primarily herbivorous crab species. Reliable morphological proxies of long-term diet strategies in herbivorous crustaceans therefore remain elusive.

## Acknowledgements

This work was supported by Brigham Young University.

